# A Structure-Aware Generative AI Framework for Revealing Functional Relationships in Proteins Families

**DOI:** 10.1101/2025.09.18.676787

**Authors:** Divyanshu Shukla, Jonathan Martin, Faruck Morcos, Davit A. Potoyan

## Abstract

The rapid expansion of protein sequence databases has far outpaced experimental structure determination, leaving many unannotated sequences, particularly the more remote homologs with low sequence identity. Because protein folds are more conserved and functionally informative than sequences alone, structural information offers a powerful lens for analysis. Here, we introduce a generative, structure-aware framework that integrates geometric encoding and coevolutionary constraints to map, cluster, and design protein sequences. Our approach employs the 3D interaction (3Di) alphabet to convert local residue geometries into compact, 20-state discrete representations. Using ProstT5, we enable bidirectional translation between amino acid sequences and 3Di representations, facilitating sensitive homology detection and structure-guided sequence generation. We then augment the latent generative landscape methodology by combining 3Di-based alignments with direct coupling analysis (DCA) and variational autoencoders (VAE), imbuing tasks such as clustering, annotation, and design with structural information. This integrative framework enhances the detection of coevolutionary signals and enables rational sampling of structural variants, even without functional labels. We demonstrate the utility of our method across diverse protein families, including globins, kinases, and malate dehydrogenases, achieving improved contact prediction, homology inference, and sequence generation. Together, our approach offers a quantitative, generative view of protein structure space, advancing protein evolution and design studies.

**Significance Statement:** Protein sequence databases are growing far faster than our ability to experimentally determine structures, leaving much of protein space poorly annotated, especially for distant homologs. Because protein structure is more conserved and informative than sequence alone, new approaches are needed to exploit structural signals at scale. We present a generative framework that integrates compact structural representations with evolutionary constraints to map, cluster, and design protein sequences. By combining geometric encoding with coevolutionary modeling, our approach enables sensitive homology detection, improved inference of structural contacts, and rational exploration of sequence space without requiring functional labels. This work provides a quantitative bridge between protein sequence and structure, advancing our ability to interpret protein evolution and guide protein design.

## Introduction

Proteins underpin a vast array of cellular functions, from energy metabolism to cell division. Deciphering their three-dimensional (3D) structures is key to understanding function, tracing evolutionary relationships, and guiding drug design^1–3^. While protein sequence databases now contain hundreds of millions of entries, experimentally determined structures remain limited due to the time and cost of traditional methods. Recent advances in computational prediction—most notably AlphaFold2 have revolutionized structural biology by providing high-accuracy models at scale^4,5^. These models now support diverse applications, including structural alignment, pocket detection, complex modeling, novel fold discovery, and genome annotation refinement^6–8^. Despite the success of sequence-based tools, remote homology detection remains a major bottleneck, leaving a substantial fraction of proteins functionally unannotated^9,10^. It is well understood that the sequence space is larger than the fold space with multiple sequences encoding the same fold, even below 30% identity. Structure-based analysis provides a powerful alternative for identifying distant homologs and revealing functional and evolutionary relationships^9,11^.

The 3D interaction (3Di) alphabet encodes the spatial relationship between each residue and its nearest neighbor into one of 20 discrete geometric states, enabling scalable structure-based comparisons. Compared to classical structural alphabets, 3Di offers lower sequential dependency, more balanced state distributions, and higher information density localized within conserved structural cores^1,12^. ProstT5 builds upon this by fine-tuning the ProtT5 language model to translate bidirectionally between amino acid and 3Di sequences^13^. This enables sensitive remote homology detection, rapid structure-aware searches, and de novo sequence generation from structural input^14,15^. Here we extend this capability by combining the 3Di representations of sequences with the latent generative landscape (LGL) model, a method previously shown to cluster sequences based on functional and phylogenetic relationships and delineate between these clusters using the energy function from Direct Coupling Analysis^17^. By using 3Di sequences instead of amino acid sequences, we can study the structural clustering of sequences to understand relationships between remotely homologous sequences and study the functional relationships between sequences with a stronger emphasis on structural features and connectivities^18^.

We demonstrate the utility of combining 3Di sequences with the LGL across several families, showcasing how the 3Di encoding transforms the information content of sequences, enabling structure-focused functional feature identification, remote homology detection, and structurally-derived phylogenetic relationships. Together, these advances yield a new quantitative and conceptual map of protein structure space, offering insights into the structural side of fitness landscapes, evolutionary diversification, and protein design.

## Results

### Structure-Informed Landscapes captures Structural Clustering and Functional Annotations

We generated structure-informed sequence landscapes by integrating ProstT5, 3Di encoding, variational autoencoders (VAE), and direct coupling analysis (DCA) (Fig. 1). Multiple sequence alignments (MSAs) for each protein family were translated into 3Di tokens by ProstT5, encoding local residue geometry. The 3Di MSAs were processed by a VAE, which mapped sequences to latent coordinates and enabled sampling across the latent space. Sequences decoded at each point were scored using a DCA-derived Potts model Hamiltonian, producing an energy landscape where low-Hamiltonian regions correspond to structurally and functionally similar sequences.

**Fig 1:**
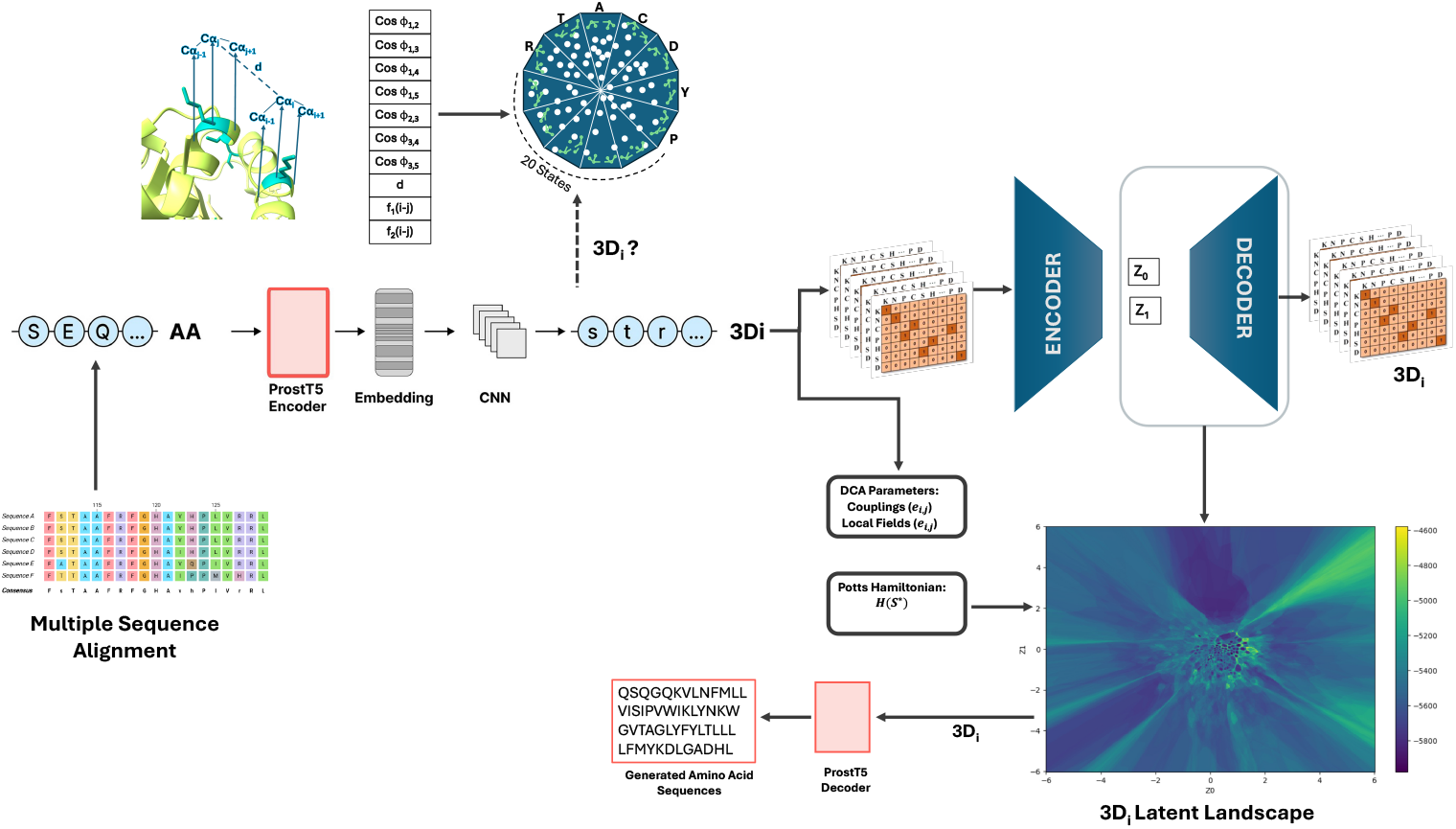
Overview of the ProstT5–3Di and VAE-based structural landscape generation pipeline. Multiple sequence alignments (MSAs) for various protein families are input into ProstT5, a pretrained language model that translates amino acid sequences into 3Di tokens representing local structural environments. MSA of these 3Di sequences are then processed through a variational autoencoder (VAE): the encoder maps sequences to latent coordinates (μ), while the decoder reconstructs them from this space. A grid of latent coordinates is sampled to generate maximum-likelihood sequences at each point. In parallel, Direct Coupling Analysis (DCA) is performed on the 3Di MSA to extract coevolutionary couplings and local fields, forming a Potts model Hamiltonian. This Hamiltonian score is used to assign an energy-based score to each decoded sequence, producing a 3D structural landscape.

Across multiple training sets, we observed that the structural landscape effectively clusters 3Di sequences according to functional and structural similarities. For instance, in the Globin family (Fig. 2A), cluster boundaries align closely with UniProt functional annotations ^33^. Similarly in Transmembrane Protein (TRPM) families, subdivided into four structurally similar groups—TRPM1/3, TRPM6/7, TRPM4/5, and TRPM2/8—which appear closely situated in the landscape (Fig. 2B) ^34,35^. Thus, by focusing on structural states represented by 3Di-based alignments yield landscapes where functionally and structurally related sequences naturally cluster. Importantly, low-Hamiltonian 3Di sequences translate into low-Hamiltonian amino-acid sequences (Fig. S1 and S2), which are typically associated with preserved protein function; consequently, this landscape can guide the ProstT5 decoder toward generating functional sequences.

**Fig 2:**
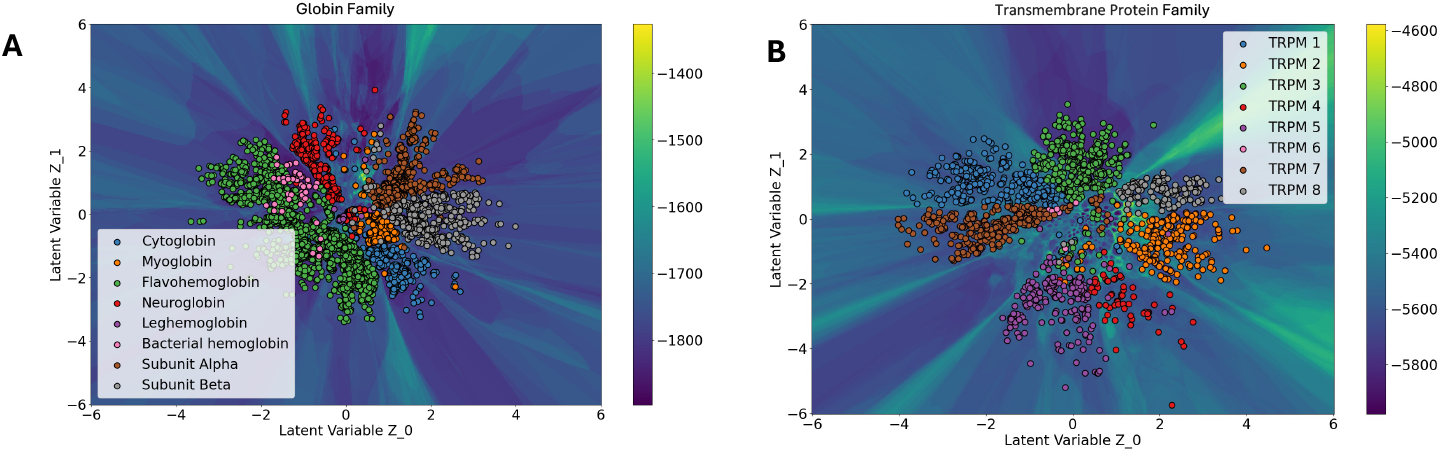
Functional and structural clustering of protein families in the 3Di-based latent landscape. (A) Globin family sequences cluster into distinct regions corresponding to UniProt-annotated functional types such as cytoglobin, myoglobin, and hemoglobin variants. (B) Transmembrane proteins (TRPM) cluster according to their known structural subdivisions—TRPM1/3, TRPM6/7, TRPM4/5, and TRPM2/8—indicating that the 3Di landscape captures architecture-driven relationships.

### Comparative Analysis of Structural and Sequence Landscapes

To assess how these landscapes capture structural similarity, we compared a conventional amino-acid (AA) MSA (Fig. 3A) with a 3Di MSA (Fig. 3B) for the malate dehydrogenase (MDH) family. In cytosolic MDHs from marine mollusks, psychrophiles and thermophiles occupy distinct regions of the AA latent space yet are known to be similar in structure, function, and dynamics^36^. We therefore sampled two cohorts from the latent space: 50 sequences from a psychrophilic region (Cluster 1) and 40 from a thermophilic region (Cluster 2). For all sequences, we predicted structures with AlphaFold^5^ and quantified pairwise similarities using Needleman–Wunsch for sequence identity and TM-score for structure^37,38^.

**Fig 3:**
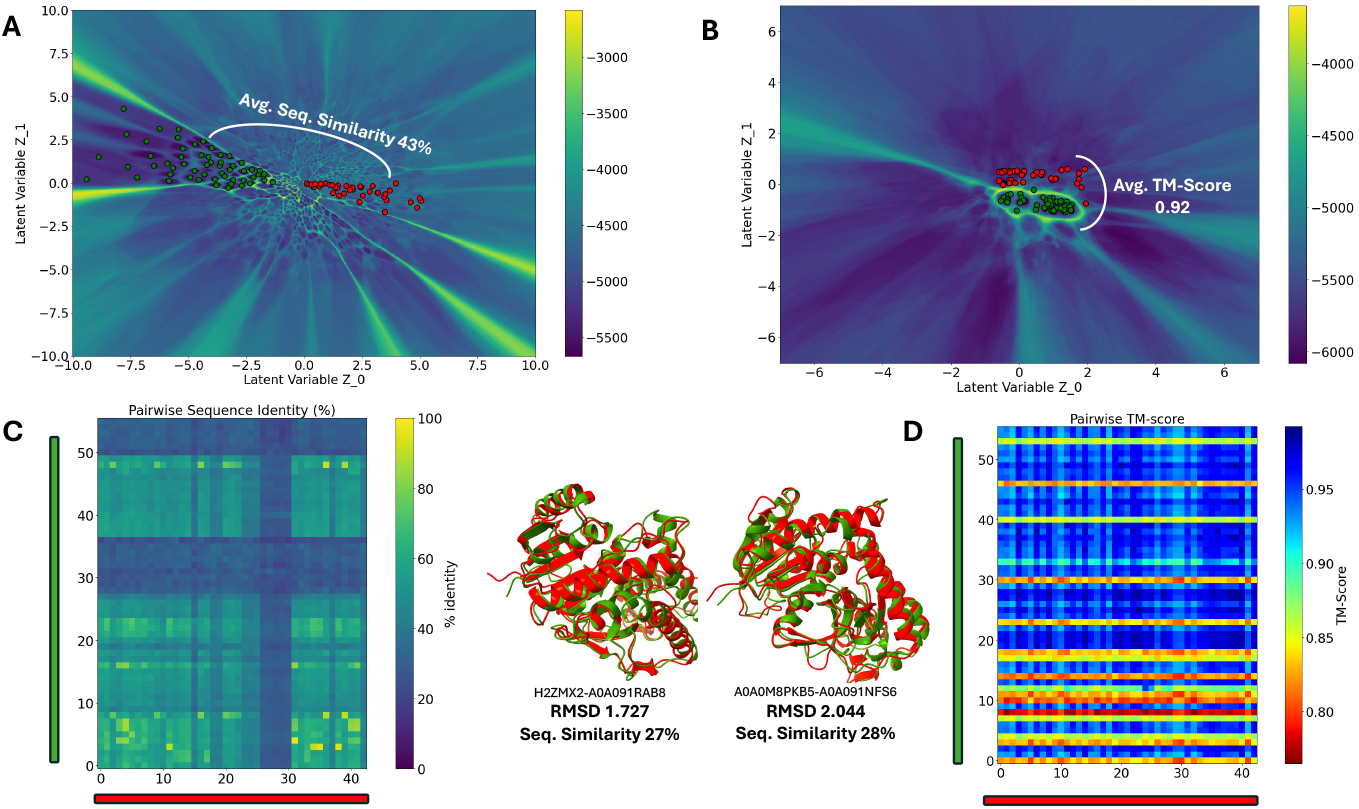
3Di reveals structural conservation across divergent MDH sequences. (A) AA-MSA latent space with two cohorts (Cluster 1: n=50; Cluster 2: n=40). (B) 3Di-MSA latent space compacts structures; cross-cluster mean TM-score ≈ 0.92. (C) Pairwise sequence identity is modest (mean ≈ 43%); example cross-cluster superpositions show low RMSDs (1.727 Å, 2.044 Å) at 27–28% identity. (D) Pairwise TM-score matrix confirms high structural similarity. 3Di reorganizes sequences by structure while Hamiltonian barriers (functional partitions) persist.

Across both cohorts, sequences share modest similarity (mean pairwise identity ≈43%; Fig. 3C), consistent with their separation in the AA landscape. In sharp contrast, their structures are highly conserved: the cross-cluster mean TM-score is ≈0.92 (Fig. 3B, 3D), well above the conventional homology threshold of 0.8^39^, and some example superpositions of structures from 2 clusters show low RMSDs with only ~27–28% sequence identity (center panels). Thus, the 3Di landscape reorganizes the sequences by structural likeness while the Hamiltonian barriers persist, demarcating putative functional partitions that were evident in the AA landscape. Together, these results show that structural constraints are strongly conserved across divergent MDH sequences and that the 3Di representation uncovers this conservation without erasing function-relevant boundaries ^40,41^.

### Decoder–Sequence Relationship and Entropy Considerations

To further investigate how encoded sequences relate to the decoder-produced landscapes, we embedded training sequences into both the sequence- and structure-based latent spaces and decoded the maximum-probability sequence at each encoded μ-coordinate. We then compared each decoded sequence’s Hamiltonian to that of its original input. In the TRPM family (Fig. 4A), both landscapes displayed a positive correlation, though the structural landscape exhibited a much stronger correlation (R=0.86) than the sequence landscape (R=0.23). This could be because the 3Di latent space has a smaller high-entropy region and Fig. 4C shows that, in both latent spaces, low-entropy regions correlate more strongly with the input sequences’ Hamiltonian than do high-entropy regions^19,42^.

**Fig 4:**
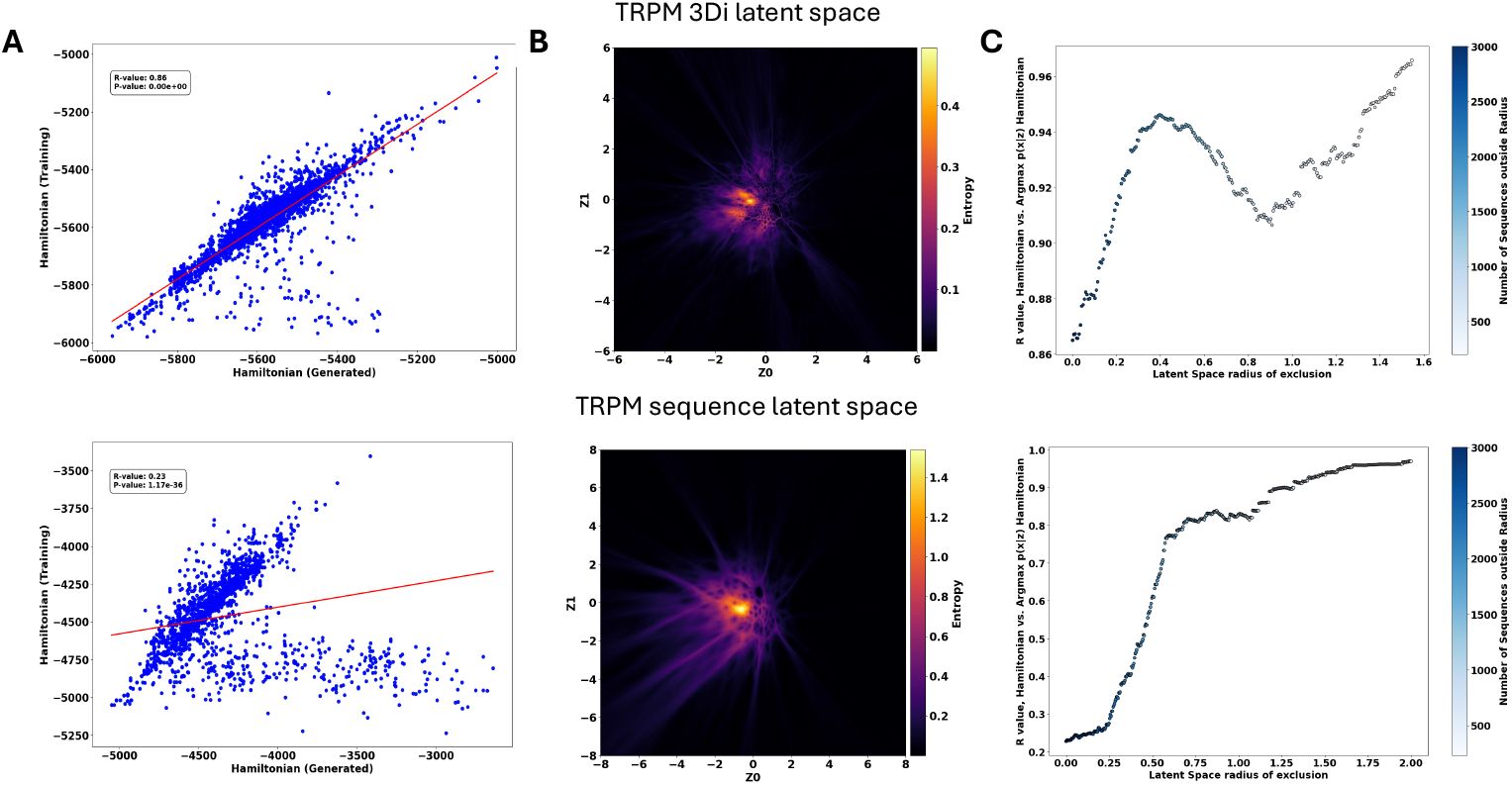
Decoder performance and entropy analysis across latent spaces for the TRPM family. (A) Comparison of DCA based Hamiltonian scores between input training sequences and sequences generated at the corresponding encoded μ-coordinate in both 3Di (top) and amino acid (bottom) landscapes. (B) Entropy of the decoder’s output distribution at each latent coordinate, calculated using Equation (11). The 3Di landscape shows lower and more localized entropy compared to the broader, higher entropy in the amino acid latent space. (C) Correlation between input and generated Hamiltonians as a function of radial exclusion from the high-entropy latent space center. Excluding central, high-uncertainty sequences improves correlation in both landscapes, with the 3Di space achieving higher correlation even near the center. Color bar indicates the number of sequences remaining after each exclusion step.

We also found that the correlation between input and generated Hamiltonians depends on the decoder’s distributional variability, which peaks at the latent-space center (Fig. 4B). Entropy (Equation 8) is highest in these regions, reflecting greater uncertainty in the decoder’s output. By excluding sequences located near these high-entropy coordinates, we consistently improved the correlation of LGL-generated (decoded) sequences (Fig. 4C). Crucially, the structural landscape retained lower entropy than its AA-based counterpart, reinforcing the notion that 3Di MSAs contain less sequence variability which smooths out the entropy of the VAE’s latent space. Although correlation levels varied by protein family (Fig. S3), all tested families showed that moving outward from the high-entropy center markedly improved the correlation between input and generated Hamiltonians. Additionally, the structural landscape typically has higher correlation in high-entropy regions; however, correlations in both landscapes converged to similar levels as the exclusion radius increased, indicating that sequences beyond the central “high-entropy” region are comparably well reconstructed in either landscape.

### Evaluating Structural Contact Prediction with 3Di-DCA and Amino Acid MSAs

To investigate the effectiveness of 3Di representations in residue-residue contact prediction, we conducted a comparative analysis of Direct Coupling Analysis (DCA) performed on 3Di-based and amino acid (AA) multiple sequence alignments (MSAs) for the globin and cysteine peptidase protein families. In the left column of Figure 5, we measure how well the direct information (DI) pairs predict PDB derived residue contacts through a True Positive Rate (TPR) over a range of the top ranked DI pairs. The top DI pairs have been well established to be predictive of residue-residue interaction in PDB structures^15^, and in all examples, we see reasonable to excellent predictive power. In the case of globin, the 3Di sequence MSA clearly outperforms the original amino acid MSA, but this was not the case in the peptidase family. By looking at the contact maps themselves, we see that in the case of globin amino acid sequences there are substantially more false positives than in the 3Di sequences. While it is true that DI pairs can find conformational changes or allosteric effects^43,44^, we could not link the false positives to any documented result, and we may assume it to be noise.

**Fig 5:**
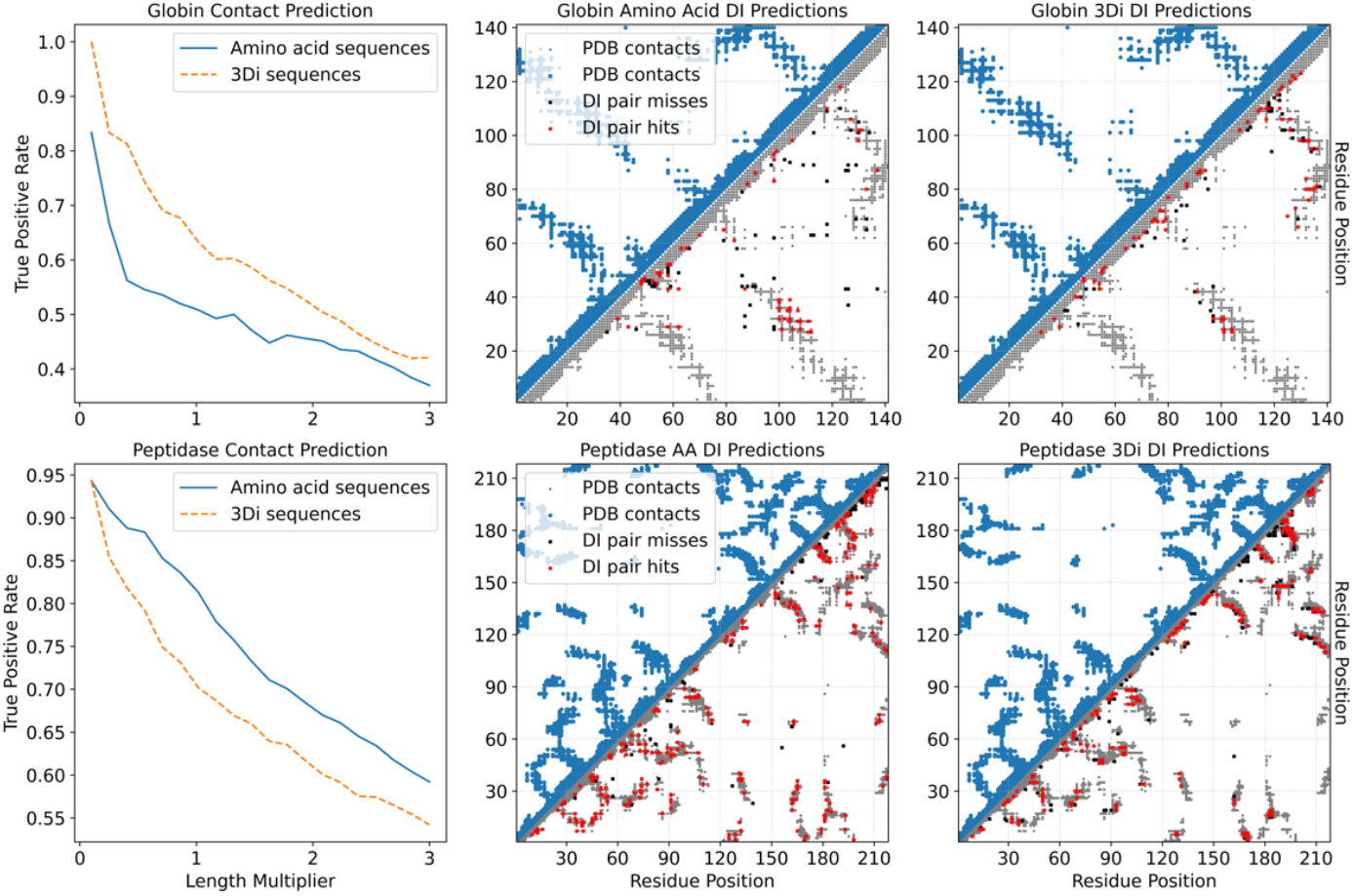
Comparison of contact predictions using 3Di and amino acid (AA) DCA respectively. (left column) Precision of long-range contacts (|i–j| > 4) across top DI pairs (0.1L to 3L). (right columns) Top L direct interaction (DI) pairs for globin (top) and peptidase (bottom), long-range predictions (|i–j| > 4). Red: true positives; Black: false positives; gray/blue: PDB derived contacts (8 Å cutoff, all-atom). Globin PDB: 1xz2. Peptidase PDB: 1AEC.

The opposite result in the peptidase family may be due to the sample size, in that the peptidase MSA has 25,531 sequences available while the globin MSA only has 10,381. We reason that with a high number of diverse real sequences in the peptidase MSA, we have better estimates of the couplings which the 3Di may be damping and thus get less true positives. This also suggests that 3Di MSA might perform better with less data, because each sequence contributes information to a fewer number of *e_ij* coupling terms due to the reduced sequence entropy (Figure 4), and this information is more directly relevant to known structures. To test this, we subsampled the globin and peptidase MSA and assessed contact prediction with smaller amounts of sequence data (Fig. S4). We see that in general 3Di sequences provide better true positive rates compared to amino acid sequences especially at lower sequence counts, and this effect is even more pronounced in the globin family.

### 3Di-based energy landscape barriers divide homologues based on dynamics similarity

Proteins that share similar folds and topological architecture often exhibit comparable dynamic behavior^45^. Since the 3Di-based landscape clusters sequences according to structural similarity, we investigated whether this clustering also captures dynamical similarity across members of the same family. To explore this, we selected representative Flavohemoglobins from distinct 3Di clusters separated by DCA based Hamiltonian barriers in the latent space (Fig. 6A). For each representative, 3D structures were predicted using AlphaFold2 and subjected to 600 ns all-atom molecular dynamics simulations. The root mean square fluctuation (RMSF) profiles of residues were then compared between pairs of proteins. We observed that proteins grouped within the same 3Di-defined cluster exhibited highly similar RMSF profiles, indicating shared dynamic behavior (Fig. 6B). In contrast, sequences from different clusters, even though all belonged to the same globin family, showed distinguishable fluctuations.

**Fig 6:**
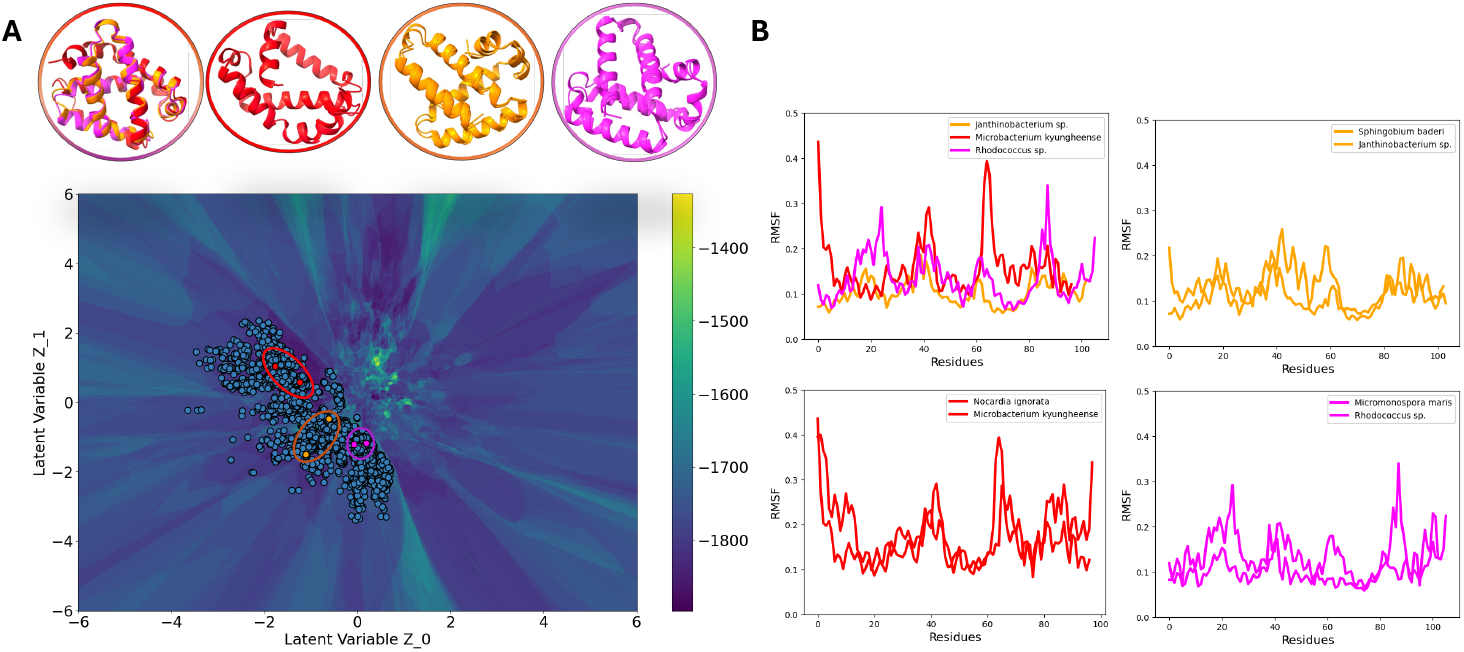
Dynamical similarity of flavohemoglobins reflected in 3Di-based latent landscape clustering. (A) 3Di latent landscape of globin family sequences colored by DCA based Hamiltonian score, with representative flavohemoglobins selected from three distinct clusters separated by Hamiltonian barriers (highlighted in red, orange, and purple). (B) Root mean square fluctuation (RMSF) profiles from 600 ns all-atom molecular dynamics simulations for AlphaFold-predicted structures of selected sequences. RMSF traces from proteins within the same cluster (e.g., red–purple, red–red) are nearly identical, indicating shared flexibility and dynamics. In contrast, proteins from separate clusters (e.g., orange–purple) show clear differences in RMSF, despite all being members of the same family.

### Evolutionary insights from 3Di landscapes of Flaviviridae glycoproteins

The Flaviviridae^46^ is a highly diverse family of enveloped positive-sense RNA viruses that includes important pathogens of humans and other animals, as well as many viruses that pose emerging threats to human health^47–49^. Previous reconstructions of Flaviviridae evolution have relied mainly on highly conserved proteins such as RNA-dependent RNA polymerase (RdRp). RdRp phylogeny supported the division of the Flaviviridae into three distinct clades: (1) an Orthoflavivirus/jingmenvirus group (2) a clade comprising the large genome flaviviruses and members of the genus Pestivirus and (3) a Pegivirus/Hepacivirus clade (Fig. S5)^50,51^. These approaches provide limited resolution of glycoproteins, which are critical determinants of viral entry, host range, and immune recognition, due to high levels of sequence divergence for example E1 glycoprotein share only 10–15% amino acid sequence identity and E2 glycoprotein sequence identity ranged from 8.5 to 15%^52,53^. E1 and E2 glycoproteins of hepaciviruses, pegiviruses, and pestiviruses form a distinct class of fusion machinery and are strictly associated with vertebrate hosts^54,55^. Using 3Di sequences of structurally aligned E1 and E2 glycoproteins, we mapped their distribution in latent space and compared them with phylogenetic trees derived from the same proteins (Fig. 7). In the E1 (structurally conseved) landscape (Fig. 7A), pegiviruses and hepaciviruses strongly overlap, reflecting structural similarity may be due to shared reliance on E1/E2 glycoproteins with internal ribosome entry site (IRES)–dependent translation^54^. By contrast, pestiviruses cluster in a distinct region, suggesting different structure orientation because of their separate evolutionary origin from an ancestor that had E glycoprotein and MTase, enabling cap-dependent translation^56^. The E2 (structurally divergent) landscape (Fig. 7B) further underscores this division: pegiviruses and hepaciviruses remain closely positioned, while pestiviruses remain divergent.

**Fig 7:**
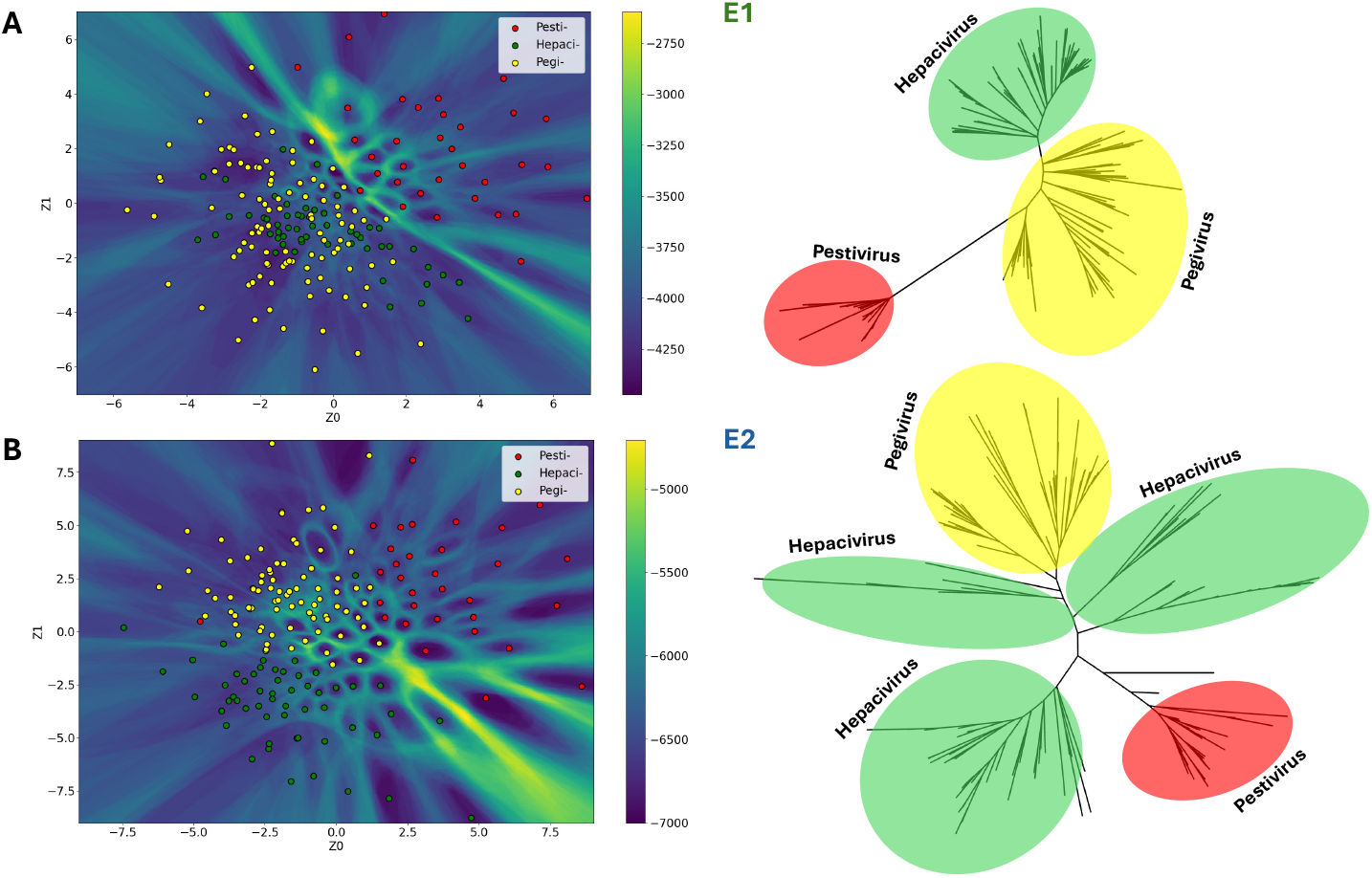
(A). Left, clustering of 3Di sequences in landscape of E1 glycoprotein of the hepaciviruses, pegiviruses and pestiviruses. Right, combined 3Di and amino acid-based E1 glycoprotein structural phylogeny. (B). Left, clustering of 3Di sequences in landscape of E2 glycoprotein of the hepaciviruses, pegiviruses and pestiviruses. Right, combined 3Di and amino acid-based E2 glycoprotein structural phylogeny.

The corresponding phylogenies (Fig. 7, right) mirror these latent-space relationships, with pegiviruses and hepaciviruses forming tight clusters and pestiviruses branching separately. Together with RdRp phylogeny, these findings align with the theory that the gain of E1E2 evolution likely arose twice independently within the Flaviviridae: once in the Pegivirus/Hepacivirus clade and once in the Pestivirus lineage^54^. This, evolutionary grouping of these proteins cannot be inferred from primary sequence due to large sequence divergence and is only revealed by structural analysis^55^.

## Discussion

Our work presents a unified, structure-aware framework for analyzing and generating protein sequences by integrating 3Di geometric encoding, variational autoencoders, and direct coupling analysis. By leveraging structural information through 3Di sequences, we construct latent generative landscapes that effectively organize proteins according to shared structural and functional features. Across diverse families, including globins, glycoproteins, malate dehydrogenases, and RNA-dependent RNA polymerase proteins, these landscapes reveal clear clustering patterns that align with known functional annotations, even for remote homologs with low sequence identity.

A key advance of our approach lies in coupling structure-informed latent spaces with coevolutionary constraints. The 3Di-based multiple sequence alignments improve latent space clustering and enhance contact prediction accuracy, particularly in families where structural conservation outpaces sequence conservation. Our analysis of decoder entropy further highlights that 3Di landscapes reduce latent space noise, supporting more reliable sequence reconstruction and generative design. Importantly, we demonstrate that sequences generated from low-Hamiltonian regions of the structural landscape retain functional domain annotations, illustrating the potential of this framework for guided sequence design.

While our findings underscore the value of integrating structural representations into generative protein modeling, several limitations remain. The 3Di alphabet, while compact and informative, abstracts local geometry and may miss longer-range structural dependencies that contribute to the function. Similarly, the latent landscapes, though effective for clustering and design, depend on the quality of input MSAs and structural predictions. Incorporating explicit functional labels, experimental binding data, or dynamics information could further enhance the model’s discriminative and generative power.

The framework presented here synergizes and complements our previous developments in latent generative landscapes using amino acid representations and opens avenues for exploring protein fitness landscapes, evolutionary trajectories, and the rational design of novel proteins with desired structural and functional properties. Extensions could include conditioning generative models on specific functional features, integrating physics-based scoring functions, or combining them with molecular dynamics to assess the stability and dynamics of generated sequences. Our work provides a foundation for structure-aware, data-driven protein modeling that bridges sequence, structure, and function in a unified generative paradigm.

## Materials and Methods

### MSA acquisition and 3Di sequence generation

For the analysis, multiple sequence alignments (MSAs) were sourced from various origins. The MSA for malate dehydrogenase was obtained using the *HMMSearch* against the UniProt database, utilizing GREMLIN ^19,20^. MSAs for globins and TRPM domains were referenced from training dataset of LGL-VAE^17^, and kinases MSAs from ^21,22^. All alignments were saved in FASTA format. To translate these amino acid sequences into 3Di sequences, we employed ProstT5—a state-of-the-art, pre-trained protein language model—on three Nvidia A100 GPUs. ProstT5, an extension of ProtT5, encodes both sequence and structural information into 3Di tokens. The generated 3Di sequences were then reformatted to align with their respective amino acid MSAs, ensuring consistency for subsequent variational autoencoder analyses^12,14^.

### VAE model architecture

The variational autoencoder (VAE) is employed to generate data samples using a latent variable(x ∈ X) model with parameters θ. This model defines a prior distribution over latent variables z, denoted as p(z). The marginal likelihood of an observed data point *x* under the model is expressed as:

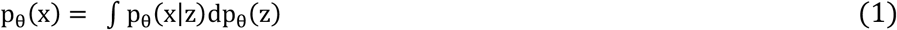

However, since both the model parameters θ and the latent variables *z* are unknown, directly evaluating the marginal likelihood (Equation 1) becomes computationally intractable—particularly as the number of parameters increases. To address this challenge, the approach introduced in ^16^ proposes approximating the true posterior distribution p(z ∣ x) with a separate, tractable distribution q_ϕ_ (z ∣ x), parameterized by *ϕ*:

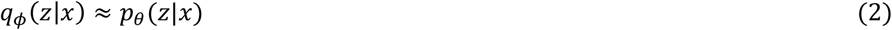

The model parameterized by ϕ, which approximates the posterior q_ϕ_ (z ∣ x), is referred to as the encoder, while the model parameterized by θ, which defines the likelihood p_θ_ (x ∣ z), is known as the decoder. Together, these models enable sampling and reconstruction of data through a latent representation.

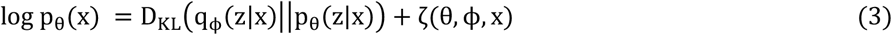

Here, D_KL_ denotes the Kullback–Leibler divergence, which measures how well the encoder’s approximate posterior q_ϕ_ (z ∣ x) aligns with the true posterior defined by the decoder. The second term in the expression represents a variational lower bound on the model’s fit to the marginal distribution over z. This leads to a simplified training objective known as the Evidence Lower Bound (ELBO), expressed as:

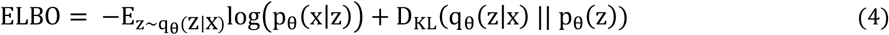

This expression defines the objective function minimized during VAE training. The first term of the ELBO represents the reconstruction loss, quantifying how accurately the decoder can reconstruct the original input from the latent representation. The second term measures how closely the learned latent distribution q_ϕ_ (z ∣ x) aligns with the assumed prior distribution p(z), using the Kullback–Leibler divergence as a regularization. To enable backpropagation through the stochastic sampling process, we apply the reparameterization trick. Specifically, the encoder is designed to output the mean μ and standard deviation σ of a Gaussian distribution. These parameters are then combined with an auxiliary noise variable ε ~ N (0, I) to produce latent variables z as:

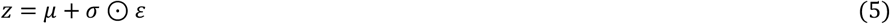

This reparametrized *z* forms the input the decoder utilizes for sequence generation, allowing us to define p_θ_ (z) as a Gaussian distribution, thereby providing an analytical solution to the gradient of Equation (4).

### Data representation and decoding

In our implementation, 3Di sequences are represented as one-hot encoded vectors. For a sequence of length L, the input array has dimensions 20 × L, where each column corresponds to one position in the sequence. Each column contains a single entry with value 1, indicating the identity of the 3Di state at that position, and 0s elsewhere. The 20 possible entries encode the 20 3Di states. During decoding, the latent variables z are passed through a Softmax activation, producing a probability distribution over the 20 possible 3Di types at each sequence position. Thus, the output layer has the same dimensions as the input: R^23×L^, where each column represents a probability distribution over sequence symbols at a given residue position.

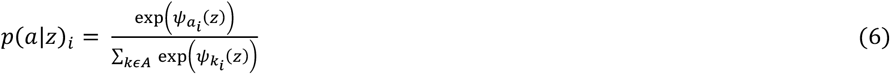

This yields L rows with probability values summing to one in each row. The reconstruction error term in Equation (4) evaluates to zero if the input and output matrices are identical, indicating that the only possible sequence at some point z is the input sequence.

### Hyperparameters and training

All models were trained using 3 × L hidden units in both the encoder and decoder, where L is the input sequence length. The ReLU activation function was applied to all hidden layers. A latent dimensionality of 2 was used. Optimization was performed using the Adam optimizer with a learning rate of 1 × 10^−4^, and L2 regularization with a penalty of 1 × 10^−4^ was applied to the hidden units. Training was terminated early if the loss failed to improve over 50 consecutive epochs. Empirically, we observed that increasing the number of hidden units beyond 3 × L did not yield improvements in validation performance for the two-dimensional latent space. All models were implemented in TensorFlow^23^ and trained either on local workstations or using NVIDIA A100 GPUs on a high-performance computing cluster.

### Landscape generation

For each trained VAE model, we constructed a Direct Coupling Analysis (DCA)^15^ model using the same input sequences used to train the VAE. The DCA model defines the probability of a sequence *S* of length *L* based on observed statistics of amino acids at individual positions A_i_ and pairs of positions (A_i_, A_j_). This probability is given by:

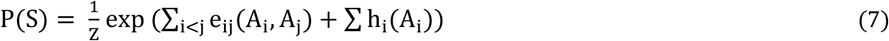

Here, e_ij_ represents pairwise coupling parameters between positions *i* and *j*, while h_i_ corresponds to a local field term that captures the frequency of amino acids at position *i*. These parameters characterize the Boltzmann-like distribution over sequences and can be inferred using various methods^24–26^.

To generate the landscape, we uniformly sampled coordinates (z_0_, z_1_) across the 2D latent space and passed them through the VAE decoder. The decoder produced a Softmax probability distribution over the 3Di states (Equation 6). From this, we extracted the maximum-probability sequence at each position as:

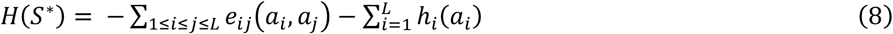

where *S*^∗^ = *a*_i_. . . . . . . *L* and 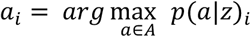

Each decoded sequence was then scored using the Hamiltonian derived from the DCA model. This Hamiltonian reflects the sequence’s likelihood under the inferred coevolutionary constraints, enabling us to map evolutionary or structural plausibility across the latent space.

### Latent space entropy calculation

To assess the uncertainty of sequence generation across the latent space, we computed the entropy landscape using the decoder’s output distribution. For each coordinate in the latent space grid, the VAE decoder produces a Softmax distribution *X* over all amino acid symbols at every residue position. The average entropy per amino acid position at each coordinate provides a measure of variability or uncertainty in the generated sequences.

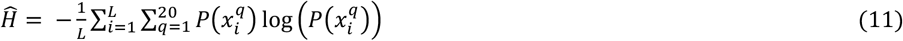

### AlphaFold2 structure prediction and performance evaluation

To evaluate the structural relevance of sequences generated by the VAE, we used Colabfold^5,27^ to predict the 3D structures of both native and decoded sequences. Representative sequences were selected from across the latent space grid and from regions of interest identified in the clustering and contact prediction analyses. All predictions were performed with no structural templates and five recycles.

The AlphaFold-generated structures were used in three core evaluations. First, to assess how well the latent space captures true structural similarity, we computed pairwise TM-scores between structures of clustered sequences. Second, to validate contact map predictions from DCA, AlphaFold structures served as reference ground truth. Residue–residue contacts were defined based on a Cβ–Cβ distance cutoff of 8 Å (Cα for glycine) and were further validated using dynamic contacts from MD simulations. Finally, we examined whether 3Di sequences with favorable DCA Hamiltonian scores also produced well-folded structures.

### Molecular dynamics simulations

The 3D structures predicted by AlphaFold2 were used as initial conformations for molecular dynamics (MD) simulations, which were performed using OpenMM^28^. Proteins were modeled using the AMBER14-all force field, and water molecules were represented using the TIP3P-FB model to construct fully solvated systems^29,30^.

Each system was first subjected to energy minimization, followed by equilibration in the NVT ensemble (constant number of particles, volume, and temperature). This was then followed by a production run in the NPT ensemble (constant pressure and temperature) for a total duration of 1.2 microseconds at 298 K. To maintain thermodynamic stability, a Langevin Middle integrator was used to regulate the system temperature at 298 K, while pressure was controlled using OpenMM’s Monte Carlo barostat to maintain 1 atm pressure^31^.

### Generation of functional sequences

To generate protein sequences with specific functional characteristics, we utilized the trained variational autoencoder (VAE) to decode latent coordinates into 3Di sequences (Fig 1). Latent coordinates were selected from regions of the landscape enriched for target domain annotations, based on their proximity to training sequences with known functions.

The decoder produced 3Di token sequences at each sampled coordinate, which were subsequently translated into amino acid sequences using the ProstT5 decoder model. To prioritize structurally conserved sequences, we selected latent coordinates based on their DCA Hamiltonian values. Low-Hamiltonian regions were favored for generating sequences hypothesized to retain native-like structure and functional domain features. The generated amino acid sequences were annotated using InterProScan^32^ to assess the presence of conserved functional domains. For structural validation, AlphaFold2 was used to predict the 3D structures of selected generated sequences, and model confidence was assessed using the per-residue *pLDDT* scores.

## Acknowledgments

This work was supported by funds from the National Institute of General Medical Sciences with grant nos. R35GM138243 (DAP) and R35GM133631 (FM). We also acknowledge support from the NIH National Institute for Allergy and Infectious Diseases with grant no. R01AI178692 (FM). FM acknowledges support from the National Science Foundation (grant no. MCB-1943442). The authors acknowledge the High Performance Computing at The University of Texas at Dallas (HPC@UTD) for providing computing resources and support.

## References

(1) Barrio-Hernandez, I.; Yeo, J.; Jänes, J.; Mirdita, M.; Gilchrist, C. L. M.; Wein, T.; Varadi, M.; Velankar, S.; Beltrao, P.; Steinegger, M. Clustering Predicted Structures at the Scale of the Known Protein Universe. Nature 2023, 622 (7983). 10.1038/s41586-023-06510-w.

(2) Chowdhury, R.; Bouatta, N.; Biswas, S.; Floristean, C.; Kharkar, A.; Roy, K.; Rochereau, C.; Ahdritz, G.; Zhang, J.; Church, G. M.; Sorger, P. K.; AlQuraishi, M. Single-Sequence Protein Structure Prediction Using a Language Model and Deep Learning. Nat Biotechnol 2022, 40 (11). 10.1038/s41587-022-01432-w.

(3) Baek, M.; DiMaio, F.; Anishchenko, I.; Dauparas, J.; Ovchinnikov, S.; Lee, G. R.; Wang, J.; Cong, Q.; Kinch, L. N.; Dustin Schaeffer, R.; Millán, C.; Park, H.; Adams, C.; Glassman, C. R.; DeGiovanni, A.; Pereira, J. H.; Rodrigues, A. V.; Van Dijk, A. A.; Ebrecht, A. C.; Opperman, D. J.; Sagmeister, T.; Buhlheller, C.; Pavkov-Keller, T.; Rathinaswamy, M. K.; Dalwadi, U.; Yip, C. K.; Burke, J. E.; Christopher Garcia, K.; Grishin, N. V.; Adams, P. D.; Read, R. J.; Baker, D. Accurate Prediction of Protein Structures and Interactions Using a Three-Track Neural Network. Science (1979) 2021, 373 (6557). 10.1126/science.abj8754.

(4) Varadi, M.; Anyango, S.; Deshpande, M.; Nair, S.; Natassia, C.; Yordanova, G.; Yuan, D.; Stroe, O.; Wood, G.; Laydon, A.; Zídek, A.; Green, T.; Tunyasuvunakool, K.; Petersen, S.; Jumper, J.; Clancy, E.; Green, R.; Vora, A.; Lutfi, M.; Figurnov, M.; Cowie, A.; Hobbs, N.; Kohli, P.; Kleywegt, G.; Birney, E.; Hassabis, D.; Velankar, S. AlphaFold Protein Structure Database: Massively Expanding the Structural Coverage of Protein-Sequence Space with High-Accuracy Models. Nucleic Acids Res 2022, 50 (D1). 10.1093/nar/gkab1061.

(5) Jumper, J.; Evans, R.; Pritzel, A.; Green, T.; Figurnov, M.; Ronneberger, O.; Tunyasuvunakool, K.; Bates, R.; Žídek, A.; Potapenko, A.; Bridgland, A.; Meyer, C.; Kohl, S. A. A.; Ballard, A. J.; Cowie, A.; Romera-Paredes, B.; Nikolov, S.; Jain, R.; Adler, J.; Back, T.; Petersen, S.; Reiman, D.; Clancy, E.; Zielinski, M.; Steinegger, M.; Pacholska, M.; Berghammer, T.; Bodenstein, S.; Silver, D.; Vinyals, O.; Senior, A. W.; Kavukcuoglu, K.; Kohli, P.; Hassabis, D. Highly Accurate Protein Structure Prediction with AlphaFold. Nature 2021, 596 (7873). 10.1038/s41586-021-03819-2.

(6) Wong, F.; Krishnan, A.; Zheng, E. J.; Stärk, H.; Manson, A. L.; Earl, A. M.; Jaakkola, T.; Collins, J. J. Benchmarking AlphaFold-enabled Molecular Docking Predictions for Antibiotic Discovery. Mol Syst Biol 2022, 18 (9). 10.15252/msb.202211081.

(7) Bordin, N.; Sillitoe, I.; Nallapareddy, V.; Rauer, C.; Lam, S. D.; Waman, V. P.; Sen, N.; Heinzinger, M.; Littmann, M.; Kim, S.; Velankar, S.; Steinegger, M.; Rost, B.; Orengo, C. AlphaFold2 Reveals Commonalities and Novelties in Protein Structure Space for 21 Model Organisms. Commun Biol 2023, 6 (1). 10.1038/s42003-023-04488-9.

(8) Sommer, M. J.; Cha, S.; Varabyou, A.; Rincon, N.; Park, S.; Minkin, I.; Pertea, M.; Steinegger, M.; Salzberg, S. L. Structure-Guided Isoform Identification for the Human Transcriptome. Elife 2022, 11. 10.7554/ELIFE.82556.

(9) Vanni, C.; Schechter, M. S.; Acinas, S. G.; Barberán, A.; Buttigieg, P. L.; Casamayor, E. O.; Delmont, T. O.; Duarte, C. M.; Eren, A. M.; Finn, R. D.; Kottmann, R.; Mitchell, A.; Sanchez, P.; Siren, K.; Steinegger, M.; Glöckner, F. O.; Fernandez-Guerra, A. Unifying the Known and Unknown Microbial Coding Sequence Space. Elife 2022, 11. 10.7554/eLife.67667.

(10) Suzek, B. E.; Wang, Y.; Huang, H.; McGarvey, P. B.; Wu, C. H. UniRef Clusters: A Comprehensive and Scalable Alternative for Improving Sequence Similarity Searches. Bioinformatics 2015, 31 (6), 926–932. 10.1093/bioinformatics/btu739.

(11) Lundin, D.; Poole, A. M.; Sjöberg, B. M.; Högbom, M. Use of Structural Phylogenetic Networks for Classification of the Ferritin-like Superfamily. Journal of Biological Chemistry 2012, 287 (24). 10.1074/jbc.M112.367458.

(12) van Kempen, M.; Kim, S. S.; Tumescheit, C.; Mirdita, M.; Lee, J.; Gilchrist, C. L. M.; Söding, J.; Steinegger, M. Fast and Accurate Protein Structure Search with Foldseek. Nat Biotechnol 2024, 42 (2). 10.1038/s41587-023-01773-0.

(13) Elnaggar, A.; Heinzinger, M.; Dallago, C.; Rehawi, G.; Wang, Y.; Jones, L.; Gibbs, T.; Feher, T.; Angerer, C.; Steinegger, M.; Bhowmik, D.; Rost, B. ProtTrans: Toward Understanding the Language of Life Through Self-Supervised Learning. IEEE Trans Pattern Anal Mach Intell 2022, 44 (10). 10.1109/TPAMI.2021.3095381.

(14) Heinzinger, M.; Weissenow, K.; Gomez Sanchez, J.; Henkel, A.; Mirdita, M.; Steinegger, M.; Rost, & B.; Heinzinger, M.; Weissenow, K.; Gomez Sanchez, J.; Henkel, A.; Mirdita, M.; Steinegger, M.; Prostt5, R. Bilingual Language Model for Protein Sequence and Structure. bioRxiv 2024.

(15) Morcos, F.; Pagnani, A.; Lunt, B.; Bertolino, A.; Marks, D. S.; Sander, C.; Zecchina, R.; Onuchic, J. N.; Hwa, T.; Weigt, M. Direct-Coupling Analysis of Residue Coevolution Captures Native Contacts across Many Protein Families. Proc Natl Acad Sci U S A 2011, 108 (49). 10.1073/pnas.1111471108.

(16) Kingma, D. P.; Welling, M. Auto-Encoding Variational Bayes. In 2nd International Conference on Learning Representations, ICLR 2014 - Conference Track Proceedings; 2014. 10.61603/ceas.v2i1.33.

(17) Ziegler, C.; Martin, J.; Sinner, C.; Morcos, F. Latent Generative Landscapes as Maps of Functional Diversity in Protein Sequence Space. Nat Commun 2023, 14 (1). 10.1038/s41467-023-37958-z.

(18) Anton, B.; Besalú, M.; Fornes, O.; Bonet, J.; Molina, A.; Molina-Fernandez, R.; De Las Cuevas, G.; Fernandez-Fuentes, N.; Oliva, B. On the Use of Direct-Coupling Analysis with a Reduced Alphabet of Amino Acids Combined with Super-Secondary Structure Motifs for Protein Fold Prediction. NAR Genom Bioinform 2021, 3 (2). 10.1093/nargab/lqab027.

(19) Lesk, A. M.; Chothia, C. How Different Amino Acid Sequences Determine Similar Protein Structures: The Structure and Evolutionary Dynamics of the Globins. J Mol Biol 1980, 136 (3). 10.1016/0022-2836(80)90373-3.

(20) Finn, R. D.; Clements, J.; Eddy, S. R. HMMER Web Server: Interactive Sequence Similarity Searching. Nucleic Acids Res 2011, 39 (SUPPL. 2). 10.1093/nar/gkr367.

(21) Kalaivani, R.; Reema, R.; Srinivasan, N. Recognition of Sites of Functional Specialisation in All Known Eukaryotic Protein Kinase Families. PLoS Comput Biol 2018, 14 (2). 10.1371/journal.pcbi.1005975.

(22) Mistry, J.; Chuguransky, S.; Williams, L.; Qureshi, M.; Salazar, G. A.; Sonnhammer, E. L. L.; Tosatto, S. C. E.; Paladin, L.; Raj, S.; Richardson, L. J.; Finn, R. D.; Bateman, A. Pfam: The Protein Families Database in 2021. Nucleic Acids Res 2021, 49 (D1). 10.1093/nar/gkaa913.

(23) Rucci, M.; Casile, A. {TensorFlow}: Large-Scale Machine Learning on Heterogeneous Systems}. Network: Computation in Neural Systems.

(24) Trinquier, J.; Uguzzoni, G.; Pagnani, A.; Zamponi, F.; Weigt, M. Efficient Generative Modeling of Protein Sequences Using Simple Autoregressive Models. Nat Commun 2021, 12 (1). 10.1038/s41467-021-25756-4.

(25) Figliuzzi, M.; Barrat-Charlaix, P.; Weigt, M. How Pairwise Coevolutionary Models Capture the Collective Residue Variability in Proteins? Mol Biol Evol 2018, 35 (4). 10.1093/molbev/msy007.

(26) Ekeberg, M.; Lövkvist, C.; Lan, Y.; Weigt, M.; Aurell, E. Improved Contact Prediction in Proteins: Using Pseudolikelihoods to Infer Potts Models. Phys Rev E Stat Nonlin Soft Matter Phys 2013, 87 (1). 10.1103/PhysRevE.87.012707.

(27) Mirdita, M.; Schütze, K.; Moriwaki, Y.; Heo, L.; Ovchinnikov, S.; Steinegger, M. ColabFold: Making Protein Folding Accessible to All. Nat Methods 2022, 19 (6). 10.1038/s41592-022-01488-1.

(28) Eastman, P.; Swails, J.; Chodera, J. D.; McGibbon, R. T.; Zhao, Y.; Beauchamp, K. A.; Wang, L. P.; Simmonett, A. C.; Harrigan, M. P.; Stern, C. D.; Wiewiora, R. P.; Brooks, B. R.; Pande, V. S. OpenMM 7: Rapid Development of High Performance Algorithms for Molecular Dynamics. PLoS Comput Biol 2017, 13 (7). 10.1371/journal.pcbi.1005659.

(29) Wang, L. P.; Martinez, T. J.; Pande, V. S. Building Force Fields: An Automatic, Systematic, and Reproducible Approach. Journal of Physical Chemistry Letters 2014, 5 (11). 10.1021/jz500737m.

(30) Jorgensen, W. L.; Chandrasekhar, J.; Madura, J. D.; Impey, R. W.; Klein, M. L. Comparison of Simple Potential Functions for Simulating Liquid Water. J Chem Phys 1983, 79 (2). 10.1063/1.445869.

(31) Martyna, G. J.; Tobias, D. J.; Klein, M. L. Constant Pressure Molecular Dynamics Algorithms. J Chem Phys 1994, 101 (5). 10.1063/1.467468.

(32) Quevillon, E.; Silventoinen, V.; Pillai, S.; Harte, N.; Mulder, N.; Apweiler, R.; Lopez, R. InterProScan: Protein Domains Identifier. Nucleic Acids Res 2005, 33 (SUPPL. 2). 10.1093/nar/gki442.

(33) Bateman, A.; Martin, M. J.; Orchard, S.; Magrane, M.; Ahmad, S.; Alpi, E.; Bowler-Barnett, E. H.; Britto, R.; Bye-A-Jee, H.; Cukura, A.; Denny, P.; Dogan, T.; Ebenezer, T. G.; Fan, J.; Garmiri, P.; da Costa Gonzales, L. J.; Hatton-Ellis, E.; Hussein, A.; Ignatchenko, A.; Insana, G.; Ishtiaq, R.; Joshi, V.; Jyothi, D.; Kandasaamy, S.; Lock, A.; Luciani, A.; Lugaric, M.; Luo, J.; Lussi, Y.; MacDougall, A.; Madeira, F.; Mahmoudy, M.; Mishra, A.; Moulang, K.; Nightingale, A.; Pundir, S.; Qi, G.; Raj, S.; Raposo, P.; Rice, D. L.; Saidi, R.; Santos, R.; Speretta, E.; Stephenson, J.; Totoo, P.; Turner, E.; Tyagi, N.; Vasudev, P.; Warner, K.; Watkins, X.; Zaru, R.; Zellner, H.; Bridge, A. J.; Aimo, L.; Argoud-Puy, G.; Auchincloss, A. H.; Axelsen, K. B.; Bansal, P.; Baratin, D.; Batista Neto, T. M.; Blatter, M. C.; Bolleman, J. T.; Boutet, E.; Breuza, L.; Gil, B. C.; Casals-Casas, C.; Echioukh, K. C.; Coudert, E.; Cuche, B.; de Castro, E.; Estreicher, A.; Famiglietti, M. L.; Feuermann, M.; Gasteiger, E.; Gaudet, P.; Gehant, S.; Gerritsen, V.; Gos, A.; Gruaz, N.; Hulo, C.; Hyka-Nouspikel, N.; Jungo, F.; Kerhornou, A.; Le Mercier, P.; Lieberherr, D.; Masson, P.; Morgat, A.; Muthukrishnan, V.; Paesano, S.; Pedruzzi, I.; Pilbout, S.; Pourcel, L.; Poux, S.; Pozzato, M.; Pruess, M.; Redaschi, N.; Rivoire, C.; Sigrist, C. J. A.; Sonesson, K.; Sundaram, S.; Wu, C. H.; Arighi, C. N.; Arminski, L.; Chen, C.; Chen, Y.; Huang, H.; Laiho, K.; McGarvey, P.; Natale, D. A.; Ross, K.; Vinayaka, C. R.; Wang, Q.; Wang, Y.; Zhang, J. UniProt: The Universal Protein Knowledgebase in 2023. Nucleic Acids Res 2023, 51 (D1). 10.1093/nar/gkac1052.

(34) Ciaglia, T.; Vestuto, V.; Bertamino, A.; González-Muñiz, R.; Gómez-Monterrey, I. On the Modulation of TRPM Channels: Current Perspectives and Anticancer Therapeutic Implications. Frontiers in Oncology. 2023. 10.3389/fonc.2022.1065935.

(35) Nilius, B.; Owsianik, G. The Transient Receptor Potential Family of Ion Channels. Genome Biology. 2011. 10.1186/gb-2011-12-3-218.

(36) Shukla, D.; Martin, J.; Morcos, F.; Potoyan, D. A. Thermal Adaptation of Cytosolic Malate Dehydrogenase Revealed by Deep Learning and Coevolutionary Analysis. J Chem Theory Comput 2025, 21 (6), 3277–3287. 10.1021/acs.jctc.4c01774.

(37) Zhang, Y.; Skolnick, J. TM-Align: A Protein Structure Alignment Algorithm Based on the TM-Score. Nucleic Acids Res 2005, 33 (7). 10.1093/nar/gki524.

(38) Needleman, S. B.; Wunsch, C. D. A General Method Applicable to the Search for Similarities in the Amino Acid Sequence of Two Proteins. J Mol Biol 1970, 48 (3). 10.1016/0022-2836(70)90057-4.

(39) Xu, J.; Zhang, Y. How Significant Is a Protein Structure Similarity with TM-Score = 0.5? Bioinformatics 2010, 26 (7). 10.1093/bioinformatics/btq066.

(40) Tian, H.; Jiang, X.; Xiao, S.; La Force, H.; Larson, E. C.; Tao, P. LAST: Latent Space-Assisted Adaptive Sampling for Protein Trajectories. J Chem Inf Model 2023, 63 (1). 10.1021/acs.jcim.2c01213.

(41) Ding, X.; Zou, Z.; Brooks, C. L. Deciphering Protein Evolution and Fitness Landscapes with Latent Space Models. Nat Commun 2019, 10 (1). 10.1038/s41467-019-13633-0.

(42) Puente-Lelievre, C.; Malik, A. J.; Douglas, J.; Ascher, D.; Baker, M.; Allison, J.; Poole, A.; Lundin, D.; Fullmer, M.; Bouckert, R.; Kim, H.; Steinegger, M.; Matzke, N. Tertiary-Interaction Characters Enable Fast, Model-Based Structural Phylogenetics beyond the Twilight Zone. bioRxiv 2024.

(43) Xie, J.; Zhang, W.; Zhu, X.; Deng, M.; Lai, L. Coevolution-Based Prediction of Key Allosteric Residues for Protein Function Regulation. Elife 2023, 12. 10.7554/eLife.81850.

(44) Morcos, F.; Jana, B.; Hwa, T.; Onuchic, J. N. Coevolutionary Signals across Protein Lineages Help Capture Multiple Protein Conformations. Proceedings of the National Academy of Sciences 2013, 110 (51), 20533–20538. 10.1073/pnas.1315625110.

(45) Keskin, O.; Jernigan, R. L.; Bahar, I. Proteins with Similar Architecture Exhibit Similar Large-Scale Dynamic Behavior. Biophys J 2000, 78 (4). 10.1016/S0006-3495(00)76756-7.

(46) Simmonds, P.; Becher, P.; Bukh, J.; Gould, E. A.; Meyers, G.; Monath, T.; Muerhoff, S.; Pletnev, A.; Rico-Hesse, R.; Smith, D. B.; Stapleton, J. T. ICTV Virus Taxonomy Profile: Flaviviridae. Journal of General Virology 2017, 98 (1). 10.1099/jgv.0.000672.

(47) Hubálek, Z.; Halouzka, J. West Nile Fever - A Reemerging Mosquito-Borne Viral Disease in Europe. Emerg Infect Dis 1999, 5 (5). 10.3201/eid0505.990505.

(48) Wang, Z.-D.; Wang, B.; Wei, F.; Han, S.-Z.; Zhang, L.; Yang, Z.-T.; Yan, Y.; Lv, X.-L.; Li, L.; Wang, S.-C.; Song, M.-X.; Zhang, H.-J.; Huang, S.-J.; Chen, J.; Huang, F.-Q.; Li, S.; Liu, H.-H.; Hong, J.; Jin, Y.-L.; Wang, W.; Zhou, J.-Y.; Liu, Q. A New Segmented Virus Associated with Human Febrile Illness in China. New England Journal of Medicine 2019, 380 (22). 10.1056/nejmoa1805068.

(49) Kartashov, M. Y.; Gladysheva, A. V.; Shvalov, A. N.; Tupota, N. L.; Chernikova, A. A.; Ternovoi, V. A.; Loktev, V. B. Novel Flavi-like Virus in Ixodid Ticks and Patients in Russia. Ticks Tick Borne Dis 2023, 14 (2). 10.1016/j.ttbdis.2022.102101.

(50) Paraskevopoulou, S.; Käfer, S.; Zirkel, F.; Donath, A.; Petersen, M.; Liu, S.; Zhou, X.; Drosten, C.; Misof, B.; Junglen, S. Viromics of Extant Insect Orders Unveil the Evolution of the Flavi-like Superfamily. Virus Evol 2021, 7 (1). 10.1093/ve/veab030.

(51) Shi, M.; Lin, X.-D.; Vasilakis, N.; Tian, J.-H.; Li, C.-X.; Chen, L.-J.; Eastwood, G.; Diao, X.-N.; Chen, M.-H.; Chen, X.; Qin, X.-C.; Widen, S. G.; Wood, T. G.; Tesh, R. B.; Xu, J.; Holmes, E. C.; Zhang, Y.-Z. Divergent Viruses Discovered in Arthropods and Vertebrates Revise the Evolutionary History of the Flaviviridae and Related Viruses. J Virol 2016, 90 (2). 10.1128/jvi.02036-15.

(52) Petrone, M. E.; Grove, J.; Mélade, J.; Mifsud, J. C. O.; Parry, R. H.; Marzinelli, E. M.; Holmes, E. C. A ~40-Kb Flavi-like Virus Does Not Encode a Known Error-Correcting Mechanism. Proceedings of the National Academy of Sciences 2024, 121 (30). 10.1073/pnas.2403805121.

(53) Garry, C. E.; Garry, R. F. Proteomics Computational Analyses Suggest That the Envelope Glycoproteins of Segmented Jingmen Flavi-like Viruses Are Class II Viral Fusion Proteins (β-Penetrenes) with Mucin-like Domains. Viruses 2020, 12 (3). 10.3390/v12030260.

(54) Mifsud, J. C. O.; Lytras, S.; Oliver, M. R.; Toon, K.; Costa, V. A.; Holmes, E. C.; Grove, J. Mapping Glycoprotein Structure Reveals Flaviviridae Evolutionary History. Nature 2024, 633 (8030), 695–703. 10.1038/s41586-024-07899-8.

(55) Oliver, M. R.; Toon, K.; Lewis, C. B.; Devlin, S.; Gifford, R. J.; Grove, J. Structures of the Hepaci-, Pegi-, and Pestiviruses Envelope Proteins Suggest a Novel Membrane Fusion Mechanism. PLoS Biol 2023, 21 (7 July). 10.1371/journal.pbio.3002174.

(56) Arhab, Y.; Bulakhov, A. G.; Pestova, T. V.; Hellen, C. U. T. Dissemination of Internal Ribosomal Entry Sites (IRES) between Viruses by Horizontal Gene Transfer. Viruses. 2020. 10.3390/v12060612.

