## Supporting Information for "A Structure-Aware Generative AI Framework for Revealing Functional Relationships in Proteins Families"

### A Structure-Aware Generative Framework for Exploring Structural and Functional Relationships in Proteins

**This PDF file includes:**

Supporting text

Figures S1 to S5

### Supporting Text

In this Supplementary Information, we provide additional methodological details, validation analyses, and applications of the 3Di-DCA generative framework introduced in the main text. The Figure S1 presents a schematic overview of the full 3Di-DCA workflow. Starting from a learned latent landscape encoding 3Di sequences, points are sampled from selected regions of the landscape, decoded into amino-acid sequences using the ProstT5 decoder, and subsequently evaluated using amino-acid–based DCA Hamiltonian scores. This pipeline enables the identification and enrichment of sequences that preserve structurally conserved and family-specific constraints.

The relationship between 3Di landscape position and amino-acid–level fitness is quantified in Figure S2. Here, 3Di points sampled from low- and high-Hamiltonian regions of the 3Di landscape are decoded with ProstT5, and the resulting amino-acid sequences are rescored using amino-acid DCA. Across both the peptidase (Fig. S2A) and globin (Fig. S2B) families, sequences originating from low-Hamiltonian 3Di regions consistently exhibit more favorable (more negative) amino-acid Hamiltonian scores. This demonstrates that the 3Di landscape acts as an effective design map, allowing one to prioritize regions enriched in family-like, structurally compatible sequences while deprioritizing high-energy regions associated with poorer fitness.

Figure S3 examines decoder performance and entropy structure across amino-acid and 3Di latent spaces for the globin family. In high-entropy regions, the amino-acid latent space shows a stronger correlation between the Hamiltonian scores of generated and training sequences than the 3Di latent space. In contrast, in low-entropy regions both landscapes correlate well with training-set DCA scores. These results indicate that the two representations capture complementary aspects of the sequence landscape, motivating their combined use for robust functional sequence generation.

The ability of 3Di-based representations to recover structural constraints is further assessed in Figure S4, which compares long-range contact prediction performance using DCA applied to amino-acid versus 3Di multiple sequence alignments across varying MSA sizes. Precision of predicted contacts ($|i–j| > 4$) is evaluated against reference crystal structures for globin (PDB: 1xz2) and peptidase (PDB: 1AEC). Across subsampled MSAs, 3Di-based DCA achieves contact prediction performance comparable to amino-acid DCA, demonstrating that compact structural alphabets retain sufficient information for accurate structural inference.

Finally, Figure S5 illustrates the application of the 3Di landscape to a conserved viral protein family. Using RNA-dependent RNA polymerase (RdRp) sequences from the Flaviviridae family, we embed the highly conserved NS5 region into a 3Di latent space. The resulting landscape (Fig. S5A) recapitulates known phylogenetic groupings, which are consistent with an independent phylogenetic reconstruction (Fig. S5B). This example demonstrates that 3Di landscapes can capture established evolutionary relationships even within conserved loci, while offering a framework that can be extended to explore more rapidly evolving regions beyond core replication machinery.

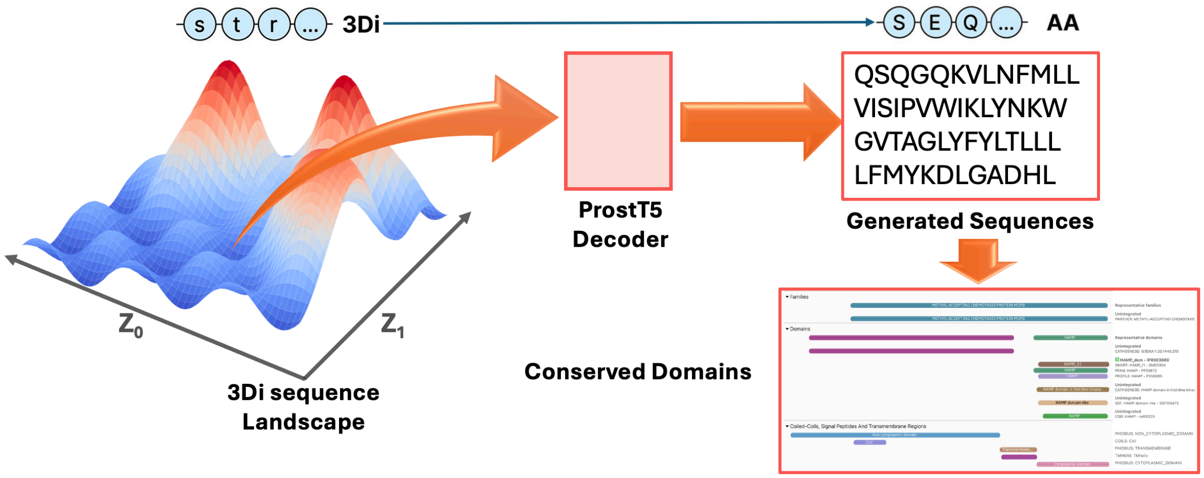


**Figure S1.** Schematic depiction of sampling from landscape encoding 3Di sequence, translating into real sequences via ProstT5 decoder and using the sequences for scoring and detecting structurally conserved domains.

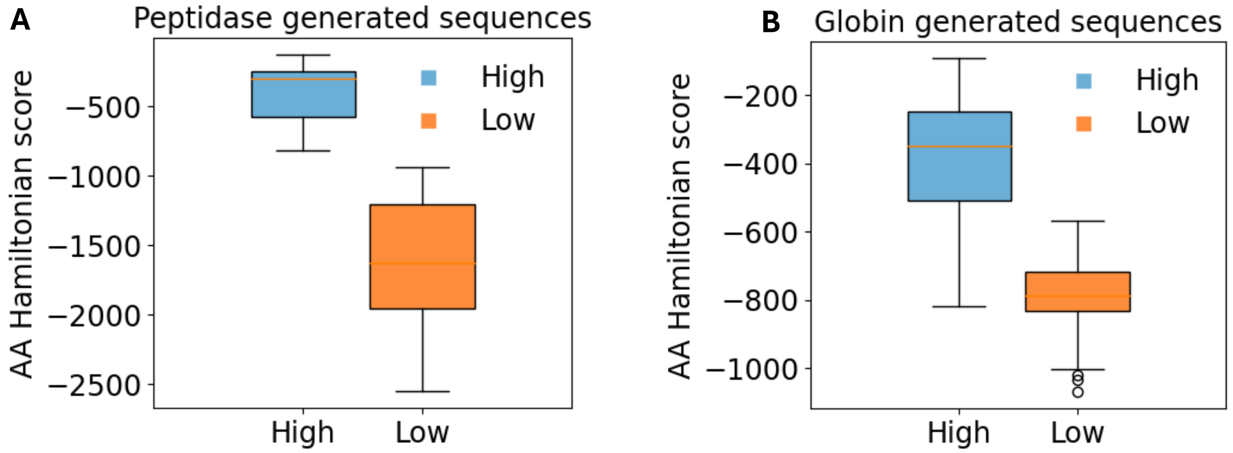


**Figure S2.** Supplementary Fig. A–B. Amino-acid Hamiltonian scores for sequences decoded by ProstT5 from 3Di seeds sampled in high (blue) vs low (orange) Hamiltonian regions of the 3Di landscape. (A) Peptidase; (B) Globin. In both families, sequences derived from low-Hamiltonian regions exhibit markedly more negative AA Hamiltonian scores than those from high-Hamiltonian regions.

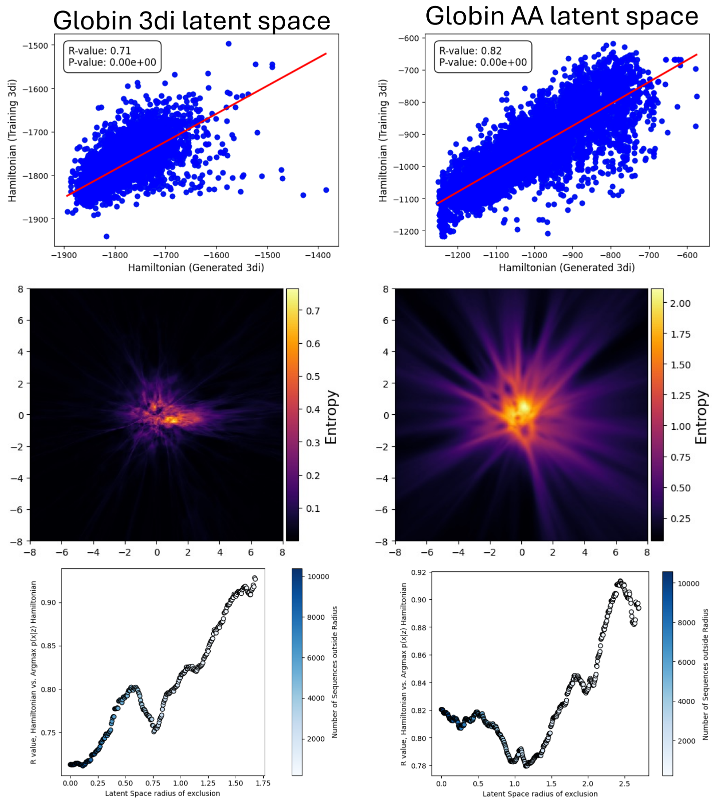


**Figure S3.** Decoder performance and entropy analysis across latent spaces for the Globin Family.

In the globin family (Supplementary Fig. 3), the AA latent space shows a stronger correlation between the DCA (Hamiltonian) scores of generated and training sequences than the 3Di latent space. This indicates that, in high-entropy regions, 3Di performance is family-specific (as is the AA space), whereas in low-entropy regions both landscapes correlates well with the DCA scores of training sequences. Together, these results argue for using both landscapes to obtain a more complete view and to guide functional protein sequence generation.

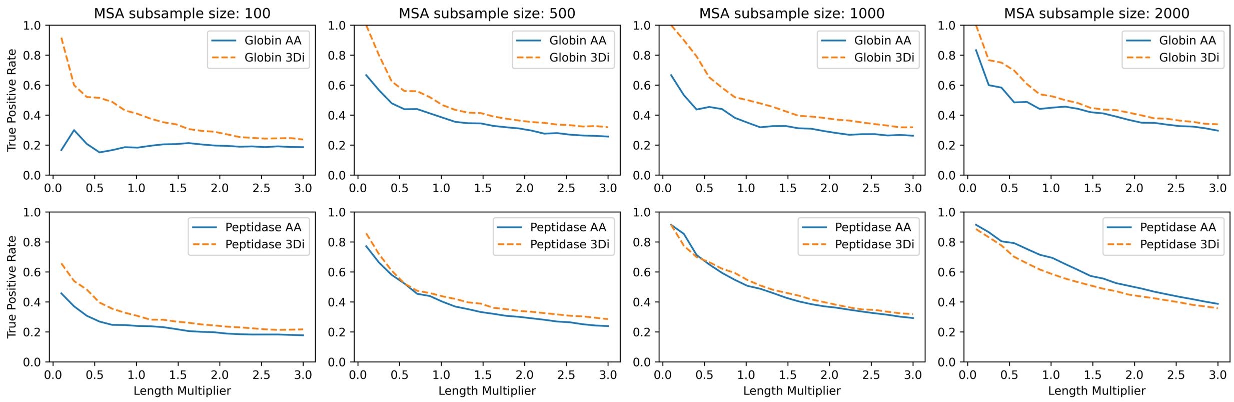


**Figure S4.** Comparison of contact predictions using 3Di and amino acid (AA) DCA across different MSA sizes. Precision of long-range contacts (|i–j| > 4) across top DI pairs (0.1L to 3L) against reference structures: Globin PDB: 1xz2. Peptidase PDB: 1AEC. Each column represents an MSA subsampled from the corresponding full MSA, measured on AA and 3Di sequences.

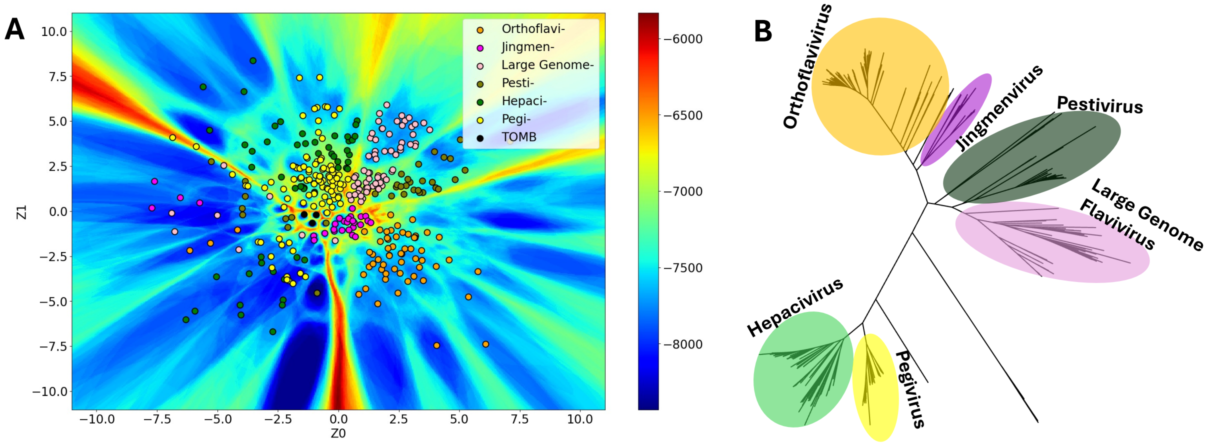


**Figure S5.** Landscape, phylogeny, and structural diversity of Flaviviridae family. (A) Latent space mapping of Flaviviridae 3Di sequences from the highly conserved NS5 region, colored by genera. (B) Phylogenetic reconstruction showing the placement of orthoflaviviruses, jingmenviruses, pestiviruses, LGF, hepaciviruses, and pegiviruses, consistent with latent space topology.
